# Early intervention with a PACAP lactoside following repetitive mild traumatic brain injury prevents persistent LHb hyperactivity and motivational deficits in male mice

**DOI:** 10.64898/2025.12.05.692609

**Authors:** Emily H. Thomas, Troy E. Smith, Lajos Z. Szabò, William J Flerlage, Fahad A. Al-Obeidi, Oana L. Rujan, Shawn C. Gouty, Regina C. Armstrong, Brian M. Cox, Torsten Falk, M. Leandro Heien, Chidiebere Ogbu, Minying Cai, Mitchell J. Bartlett, Robin Polt, Fereshteh S. Nugent

## Abstract

Mild traumatic brain injury (mTBI) often causes long-lasting mood and motivational deficits, significantly impairing quality of life. Pituitary adenylate cyclase-activating polypeptide (PACAP) exhibits neuroprotective properties, but its role in mitigating post-mTBI circuit dysfunction remains poorly understood. Using a murine repetitive closed-head mTBI model, we evaluated whether TES2320—a novel, stable, and brain-penetrant PAC1 receptor (PAC1R) agonist—could prevent or reverse post-injury alterations in the lateral habenula (LHb), a critical hub for mood and motivation.

RNAscope analysis revealed that mTBI induced a persistent, widespread reduction in PACAP mRNA alongside a compensatory increase in PAC1R mRNA within the LHb, indicating local PACAP signaling deficits. We then tested early (immediately post-injury) and late (single injection 1-month post-injury) PAC1R agonist interventions in young adult male mice. One month post-injury, both early and late interventions normalized mTBI-induced LHb spontaneous tonic hyperactivity and hyperexcitability. However, only early intervention rescued deficits in self-care grooming motivation. Conversely, late administration failed to reverse delayed grooming initiation and produced a transient reduction in total grooming behavior 24 hours post-treatment, highlighting a potential acute side effect of delayed administration.

Overall, our findings demonstrate that early PAC1R agonist intervention prevents mTBI-induced LHb hyperactivity and motivational deficits, likely by restoring persistent PACAP signaling deficits within LHb circuits. This preclinical study provides strong evidence supporting novel PAC1R agonists as a promising preventive therapy for mTBI-related affective disorders and anti-reward circuit dysfunction.

## Introduction

Mild traumatic brain injury (mTBI) is a significant global health issue, with millions affected annually. While many symptoms are transient, a major and lasting consequence is the development of chronic psychiatric conditions such as depression, anxiety, and post-traumatic stress disorder (PTSD)(1, 2). The risk for these disorders is even higher with repeated mTBIs, a serious concern for athletes, military personnel, and others in high-risk groups(3, 4). A critical challenge is that mTBI-related psychopathologies often resist standard treatments like serotonin reuptake inhibitors (SSRIs), suggesting they have different underlying causes than non-TBI mood disorders(5–7).

Endogenous neuropeptides that exploit natural repair mechanisms offer promising therapeutic avenues for addressing the cellular and molecular changes that follow brain injury. Among these, pituitary adenylate cyclase-activating polypeptide (PACAP) is a potent and versatile neuropeptide with well-established neuroprotective and neurotrophic properties(8). PACAP is a highly conserved endogenous hypothalamic neuropeptide belonging to the secretin/glucagon/vasoactive intestinal peptide (VIP) family(8). PACAP exists in two primary amidated forms, PACAP_1-27_ and PACAP_1-38_, with PACAP_1-38_ being predominant in the mammalian brain and somewhat brain penetrant. PACAP’s effects are primarily mediated via its cognate high-affinity PAC1 receptor (PAC1R), although it also interacts with VPAC1 and VPAC2 receptors(9). It is crucial for a variety of brain functions and is found in key stress-related circuits(8, 10–12). These circuits include PACAP-expressing neurons within the hypothalamus (e.g., the paraventricular nucleus of hypothalamus, PVN)(11, 12), cortical regions including medial prefrontal cortex (mPFC)(11, 13, 14), ventral hippocampus CA3(11), amygdala(15), lateral habenula (LHb)(11, 16) and brain stem (e.g., parabrachial nucleus)(11), with some subtle differences in expression strength across rodent brain regions(11). Studies show that PACAP protects tissues from damage, dampens inflammation, and inhibits apoptosis(9, 17–22), making it highly relevant to TBI. PACAP exerts these protective actions mostly through PAC1Rs, for instance, by decreasing pro-inflammatory cytokines like TNF-α, and promoting a neuroprotective microglial phenotype(22). A deficiency or dysregulation of PACAP signaling has been linked to several stress-related disorders, including depression and PTSD(23–25). Clinical and preclinical studies following TBI consistently show a reduction in PACAP expression, further supporting the idea that enhancing PACAP signaling could be a beneficial intervention(20, 26). Despite its therapeutic potential, native PACAP faces a significant clinical limitation: its rapid peripheral metabolism and poor ability to cross the blood-brain barrier (BBB)(27). To overcome this, our team developed a new class of truncated PACAP glycopeptide analogues (exemplified by PACAP_1–23_) designed for enhanced stability and BBB penetration. This design strategy was informed by prior structure-activity relationship studies showing that PACAP_1–23_ retains receptor-binding capability (9, 26–28), with further truncations confirming that the six N-terminal residues are sufficient for agonist activity (29). Although PACAP_1–23_ exhibits reduced receptor affinity compared to full-length PACAP, it retains downstream signaling and neuroprotective efficacy *in vitro*, validating its utility as an optimization scaffold. Glycosylation further improves overall pharmacokinetic profiles and BBB passage, likely by modulating amphipathicity and membrane interactions, though the precise translocation mechanism remains under investigation (9, 26, 27). Crucially, interpreting PACAP pharmacology requires caution when relying on heterologous expression systems. As Civciristov and Halls noted (30), cell-based models can yield misleading dynamic ranges for PACAP, which exerts potent physiological activity in native tissues and *in vivo* at ultra-low (picomolar to femtomolar) concentrations.

These novel glycosylated PACAP analogues have already shown promising results in other models of neurological injury. For example, they have demonstrated potent neuroprotective and anti-inflammatory effects in a fluid percussion injury model of TBI, and have also shown beneficial effects in models of Parkinson’s disease and ischemic stroke(9, 21). Specifically, a prior study showed that a single dose of the PAC1R agonist, 2LS80Mel, administered before TBI, prevented injury-induced sleep and motor deficits 7-14 days later(9). While this work established the short-term protective effects of PAC1R agonists, their potential to address the long-term emotional and motivational consequences of TBI remained unknown. Our recent studies point to the LHb as a crucial brain region involved in the negative affective symptoms seen after mTBI(31–33). The LHb acts as an “anti-reward” hub, promoting avoidance behaviors and a lack of motivation(34). In models of depression, LHb hyperactivity is a common finding associated with anhedonia and social withdrawal(34–38). Using a repetitive closed-head injury model in both male and female mice(31–33, 39), we’ve shown that mTBI causes persistent LHb hyperactivity and hyperexcitability. This hyperactivity, which appears approximately 3-4 weeks post-injury, is characterized by an increase in the synaptic excitation/inhibition (E/I) balance and heightened rebound bursting activity in the LHb. This mTBI model also produced several behavioral deficits consistent with human mTBI-related mood disorders, including, reduced motivation for self-care grooming(31, 33), altered social interactions(31, 39) and increased passive freezing behaviors(31). These findings validate our model’s translational relevance. Crucially, we established a causal link between LHb hyperactivity and motivational deficits. Using chemogenetic inhibition, we were able to reverse the self-care grooming deficits in male mice by suppressing the activity of LHb glutamatergic neurons. This demonstrates that mTBI-induced LHb hyperactivity can directly drive motivational impairments(33).

Given the extensive presence of PAC1Rs in the LHb and its downstream regions(11), along with evidence that PACAP-expressing neurons in the LHb can promote anxiolytic (anxiety-reducing) behaviors(16), we hypothesized that PACAP-PAC1R signaling might be compromised in the LHb following mTBI, contributing to the observed LHb hyperactivity and behavioral deficits. Here, we first investigated whether mTBI altered PACAP and PAC1R mRNA expression within the LHb. Subsequently, we tested whether our newly developed PAC1R agonist, TES2320(29), could prevent as well as reverse mTBI-related LHb hyperactivity and associated motivational deficits in self-care grooming behavior in male mice. Our results confirmed that mTBI does, in fact, decrease PACAP mRNA expression in the LHb. Furthermore, our most significant finding is that early intervention with our PAC1R agonist successfully prevented both the physiological (LHb hyperactivity) and behavioral (motivational) changes caused by mTBI in male mice. This provides strong evidence for the potential use of PAC1R agonists as a therapeutic strategy for treating mTBI-related reward circuit dysfunction and depressive-like behaviors.

## Materials and Methods

### Design and Synthesis of the PAC1R Agonist: TES2320

The PAC1R agonist TES2320 was developed and synthesized by the Polt Laboratory using solid-phase synthesis(9, 29). The details of the synthesis and purification of this glycopeptide, as well as its efficacy toward the PAC1 receptor in CHO cells have been previously reported(29). This compound is based on a truncated, 23-amino-acid analogue of the naturally occurring 27-amino-acid PACAP hormone (26) to which a lactoside-bearing serine-amide has been appended at the C-terminus. The lactoside moiety ensures stability and assists in brain penetration (i.e., its ability to cross the BBB). The presence of the glycoside (lactose) moiety appears to enable the analogue to activate the Class B receptor without needing to bind to its extracellular domain (Figure 1). The functional potency of TES2320 (cAMP stimulation) at the human PAC1 receptor was previously determined to be 1.8 nM in over-expressed in CHO cells (27). While these findings align with cell-based efficacy studies of PACAP and related peptide hormones, it remains important to distinguish between overall functional efficacy, initial membrane adsorption, and direct receptor binding events.

**Figure 1.**
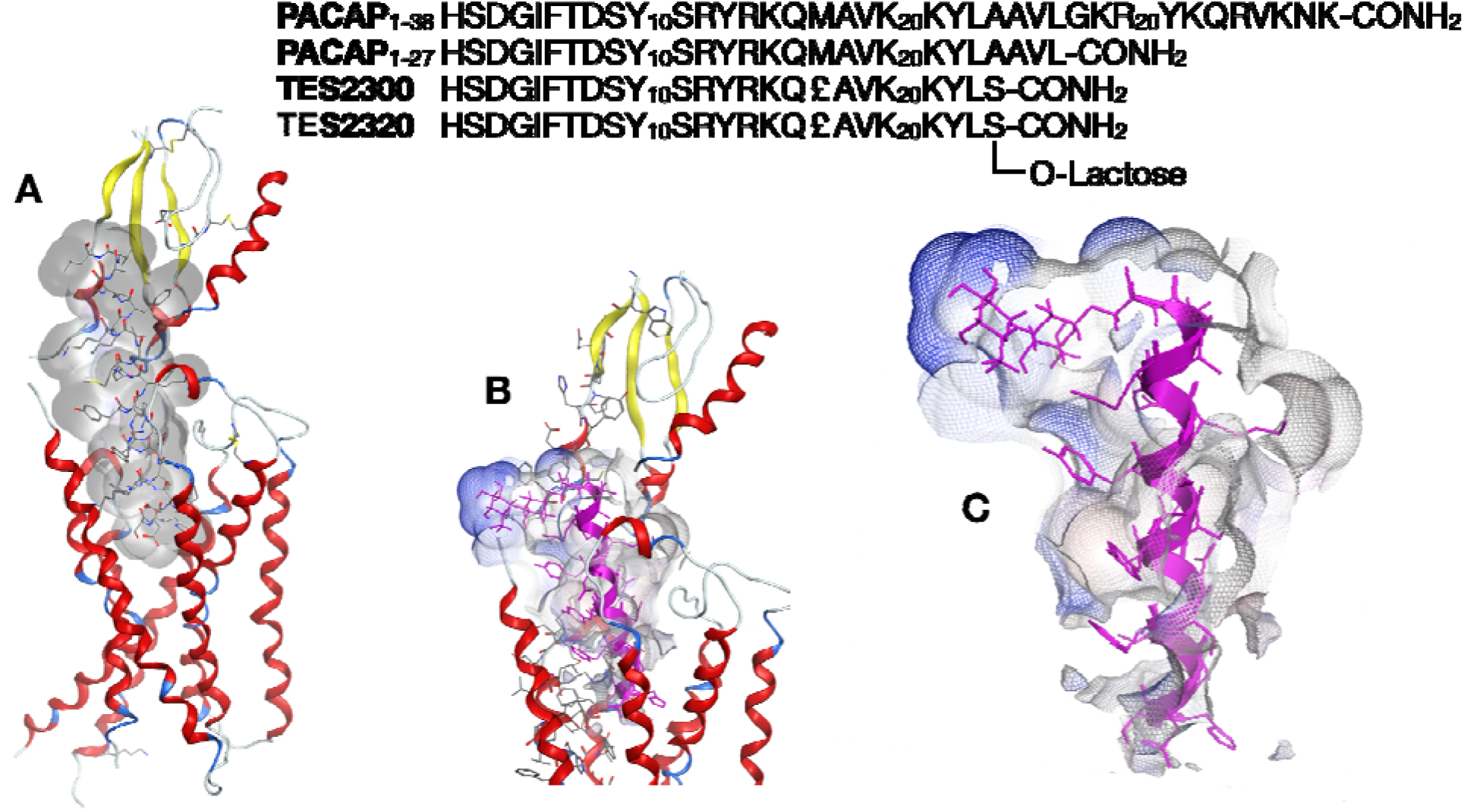
*In silico* modeling of truncated PACAP glycoside (TES2320) binding to the PAC1R. The sequences for native ligands PACAP_1–38_ and PACAP_1–27_ are displayed with the lactoside drug TES-2320. The easily oxidized residue Methionine(17) has been replaced with the non-canonical amino acid Norleucine (£). (**A**) Published cryo-electron microscopy (Cryo-EM) data of the native PACAP 27-mer bound to PAC1 (PDB: 6M1I). This structure was used as a starting point for subsequent calculations. (**B**) An *in silico* model of the glycopeptide drug TES-2320 bound to the PAC1 receptor (ADCYAP1R1) was generated with MOE^®^. (**C**) The isolated PAC1 receptor binding pocket occupied by TES-2320 is depicted. The interactions are color-coded: gray represents steric (hydrophobic) interactions and blue indicates solvent-exposed interactions. All modeling was performed using the MOE^®^ software package (Chemical Computing Group, Montreal, Canada) with the AMBER10:EHT force field, which uses specialized parameters for non-native amino acids such as serine lactoside.

Rather than relying strictly on classic ECD-mediated ligand capture, the C-terminal lactoside modification of TES2320 (and its unglycosylated counterpart TES2300) is proposed to alter membrane partitioning and initial peptide orientation at the extracellular face. Indeed, structural and biophysical studies—including nuclear magnetic resonance (NMR) analyses of isolated domains(40), highlight the structural autonomy of the ECD, suggesting that modifications along the peptide backbone can subtly alter how ligands approach the transmembrane core across PAC1, VPAC1, and VPAC2 receptors.

Our intent here is not to suggest complete independence from the ECD, but rather to highlight how C-terminal glycosylation influences peptide orientation, membrane partitioning, and receptor approach at the extracellular interface. Consistent with this rationale, our current cAMP assays effectively reproduced the published cell-based activity patterns (26). The binding assay of TES2320 towards PAC1 was measured to be 4.50 pM using cellular membrane which was highly expressed PAC1 from the CHO cells; the drug’s baseline membrane partitioning affinity (Kd) was determined to be 1.35 nM at receptor-free egg phosphatidylcholine PC membranes using Plasmon Waveguide Resonance (PWR) spectroscopy, the most sensitive technology to monitor molecular interaction in real time as previously described (29). The unglycosylated variant TES2300 showed ten fold higher affinity for PC bilayers, which was expected. The chemical formula of TES2320 is the triflate salt: HSDGIFTDSYSRYRKQ*£*AVKKYL-Ser(β-Lactose)-CONH_2_ • 2 CF_3_COOH (where *£* represents Norleucine), with a calculated molecular weight (MW) of 3372.5 g/mol.

### In Silico Structural Refinement and Binding Energy Calculations

To understand the interaction energetics of the glycopeptide **TES2320** within the human PAC1 receptor cavity, computational modeling was performed using the Molecular Operating Environment software package (MOE, Chemical Computing Group, Montreal, Canada). Because high-resolution experimental coordinates for the native PACAP•PAC1 complex are available via published Cryo-EM data(41), a stochastic *de novo* docking search was not required; the experimental coordinates were instead used to define the exact binding site and ligand orientation.

The structural coordinates of 6M1I were imported into MOE, and the bound native PACAP_1-27_ ligand was modified *in situ* to construct TES2320. Briefly, the C-terminal peptide portion (Ala^25^-Val^26^-Leu^27^) was deleted. The resulting PACAP_1-24_ intermediate was subjected to local energy minimization within the binding cavity. Subsequently, Ala^24^ was mutated to Ser(*O*-Lactose)-NH_2_ to yield the final TES2320 structure.

Hydrogen atoms were added, and optimal protonation states for all titratable residues were assigned at pH 7.4 using the *Protonate3D* protocol. Energy minimizations and structural refinements were conducted using the Amber10:EHT force field. To preserve the experimentally determined Cryo-EM backbone architecture while relieving local steric clashes introduced by the glycan modification, the refinement was performed using a rigid receptor protocol.

The placement method was set to “None” to restrict calculations to the singular, experimentally guided binding mode and to avoid the generation of misleading alternative poses stemming from peptide backbone flexibility. Binding free energy (Δ*G*) was estimated using the London dG scoring function. The single, lowest-energy refined conformation of the TES2320•PAC1 complex yielded a final binding score (*S*) of -22.16 kcal/mol with a local refinement root-mean-square deviation (RMSD) of 2.00Å.

### Plasmon Waveguide Resonance (PWR) Binding Assays

Plasmon Waveguide Resonance (PWR) spectroscopy, an enhanced technology related to Surface Plasmon Resonance (SPR) is capable of monitoring molecular anisotropy and measuring mass and structural changes in real time along with molecular interactions with receptor in the lipid bilayer. PWR was utilized to quantify label-free ligand binding affinities. Intact cell membranes containing human PAC1 receptors (purified and isolated from CHO cells, as described below) were immobilized onto the silica surface of a PWR prism element.

The prism was integrated into a PWR spectrometer system (Mainline Scientific, Malvern, PA). Stepwise cumulative injections of TES2320 were introduced into the sample reaction cell across a range of concentrations. Binding events, along with any associated conformational shifts in the receptor or lipid matrix, were directly monitored via the angular resonance shifts of parallel (*p*-polarized) and perpendicular (*s*-polarized) light.

The macroscopic dissociation constant (Kd) representing the baseline partitioning affinity of the glycopeptide into receptor-free egg phosphatidylcholine (eggPC) control membranes was determined to be 1.35 nM.

### Membrane Preparation and Receptor Isolation

Human PAC1 receptors were stably expressed in CHO cells according to previously established protocols. Cells were maintained at 37°C in a humidified 5% CO_2_ incubator in a 50:50 DMEM/F12 medium supplemented with 10% heat-inactivated fetal bovine serum (FBS), 1X penicillin-streptomycin, and 500 μg/mL G418 (neomycin) to maintain selective pressure.

For membrane isolation, cells were harvested, suspended in ice-cold membrane buffer (50 mM HEPES, pH 7.5) supplemented with 0.1% mammalian protease inhibitor cocktail (Sigma-Aldrich), and homogenized for 60 seconds using a sawtooth homogenizer. The homogenate was centrifuged at 1,500 rpm for 10 minutes at 4°C to pellet cellular debris. The supernatant was collected and subjected to ultracentrifugation at 32,750 rpm for 2 hours at 4°C.

The resulting membrane pellet was resuspended in 1.2 mL of membrane buffer, split into 300 μL aliquots, and stored at -80 C° for subsequent assays. Total membrane protein concentration was determined via a bicinchoninic acid (BCA) assay (Pierce), measuring colorimetric absorbance at 560 nm using a microplate reader (μQuant, Bio-Tek Instruments, Inc.).

#### Animals

All animal experiments adhered to the National Institutes of Health (NIH) Guide for the Care and Use of Laboratory Animals and were pre-approved by the Uniformed Services University Institutional Animal Care and Use Committee. To minimize animal suffering and the number of subjects used, we made every effort to optimize our procedures. We acquired young adult male C57BL/6 mice from the Jackson Laboratory (JAX) at approximately postnatal day 35-49 (PN35-49). Upon arrival, mice were given a minimum of 72 hours to acclimate to the vivarium. They were then group-housed in standard cages within a temperature-controlled room. The vivarium was maintained on a 12-hour light/12-hour dark cycle (lights on: 06:00-18:00) with ad libitum access to food and water. All experimental procedures were conducted within a specific timeframe: 2 to 4 hours after the start of the light cycle.

#### Carotid Artery Catheterization

Blood–brain barrier penetration and pharmacokinetics of TES2320 were determined by simultaneous, and time-locked, microdialysis and blood sampling per protocols that are well-established in our laboratory(42–46). Mice (PN56-70) were anesthetized with isoflurane (1%) mixed with medical grade oxygen (0.75 L/min). Two incisions were made: a 0.5 cm incision in between the scapulae and a 1-cm ventral neck incision right of midline. The right carotid artery was isolated, ligated cranially with 4-0 silk, and temporarily occluded at the caudal end of the vessel with a micro clamp. A small incision was made into the carotid artery and PE-10 tubing, preflushed with heparin/glycerol (500 IU/ml, 50:50) and connected to a 28-gauge PinPort^TM^ (PNP3F28; Instech, Plymouth Meeting, PA), was advanced toward the heart. The clamp was released, the caudal ligature secured the catheter, and the catheter was tunneled subcutaneously to the scapular incision. Both incisions were closed with staples. The mouse was immediately transferred to a stereotaxic frame to implant the microdialysis probe.

#### Microdialysis Surgery

A midsagittal skull incision was followed by a craniotomy (AP +0.5 mm, ML +2.0 mm from bregma). The dura was removed and a microdialysis probe (CMA12, 8309664, 4 mm membrane, 100 kDa cutoff; HBio, Holliston, MA) was inserted into the striatum (DV −5.0 mm).

#### In Vivo Microdialysis and Pharmacokinetic Analysis

Microdialysis probes were primed with artificial cerebrospinal fluid aCSF (15 mM Tris HCl, 10 mM Tris base, 126 mM NaCl, 2.5 KCl, 1.2 mM NaH_2_PO_4_, 2mM Na_2_SO_4_, and adjusted pH to 7.4) containing 0.05% bovine serum albumin (BSA) at a rate of 1 μL/minute using a two channel dual syringe pump (Fusion 101A Syringe Pump; Chemyx, Stafford, TX) for one-hour and then placed into the striatum at the coordinates listed above for 30-minutes prior to collecting baseline dialysate. Using the other pump channel, the preservation solution containing 150 ng/mL DADLE ([D-Ala2, D-Leu5]-Enkephalin; 10% acetic acid in 0.05% BSA in aCSF as an internal standard) is combined with the dialysate immediately *via* a micro-Tee at a flow rate of 1 μL/min (Figure 2A.). 3 mg/kg each (6 mg/kg total drug) of TES2300 and TES2320 were administered intraperitoneal (IP) at t = 0. Dialysate was collected at −10, +10, +20, +30, +40, +50, and +60 minutes. Blood was collected at −5, +1, +5, +15, +25, +35, +45, and +55 minutes to correspond to the midpoint of the dialysate fraction. To separate plasma from whole blood, each sample was collected in ethylenediamine tetraacetic acid capillary tubes (CT-CB300-K2EDTA; Sarstedt, Nümbrecht, Germany), inverted repeatedly for 30 seconds, and centrifuged for 5 minutes at 2,500 x g in a refrigerated centrifuge (Galaxy Mini Centrifuge; VWR, Radnor, PA).

**Figure 2.**
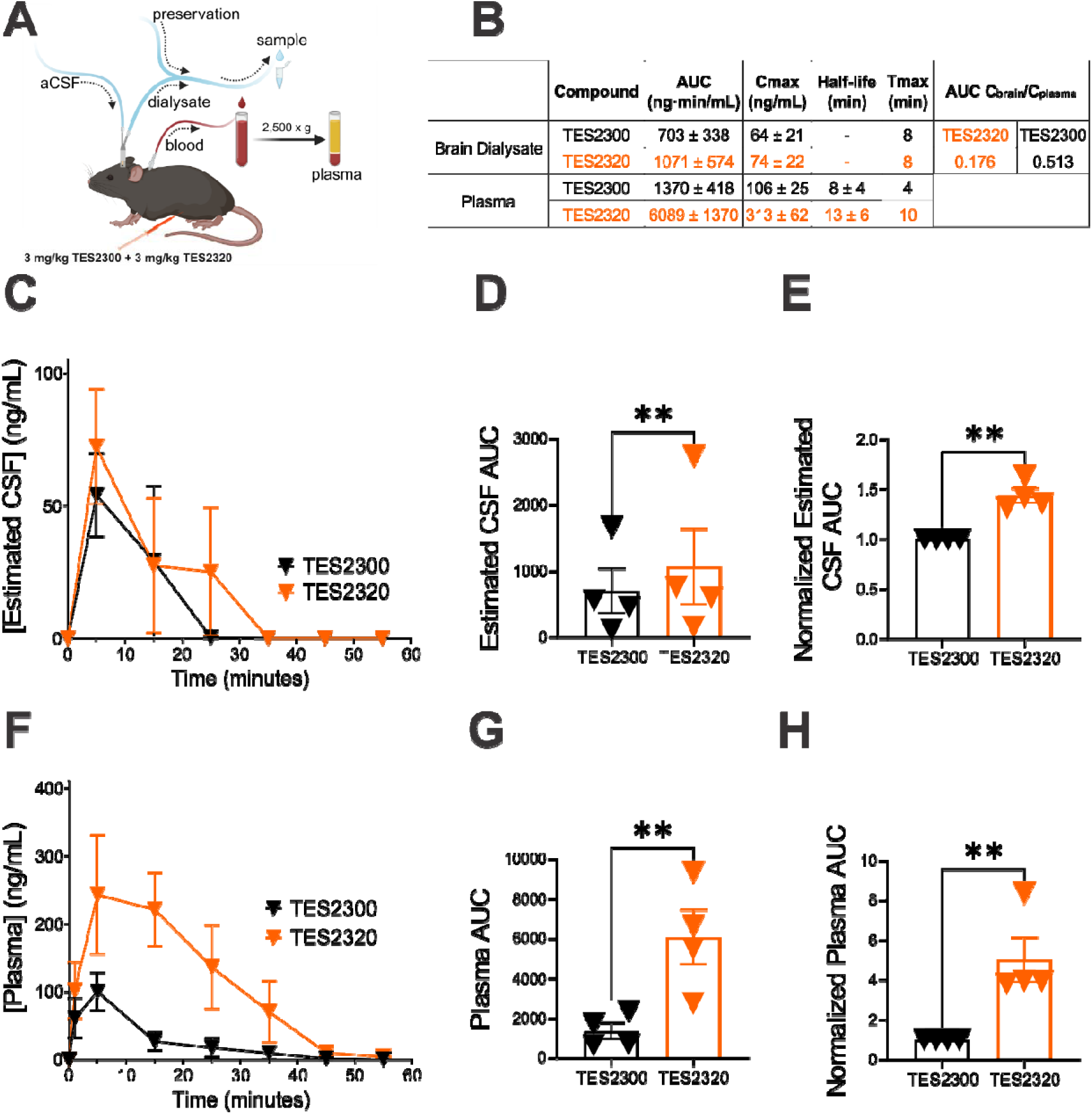
Glycosylation of PACAP improves blood-brain barrier penetration and serum half-life in C57Bl/6J mice. **(A)** Microdialysis and pharmacokinetics experimental schematic. The carotid artery was catheterized and a microdialysis probe was placed in the mouse striatum. The unglycosylated peptide amide TES2300 (3 mg/kg) and lactoside-bearing drug candidate TES2320 (3 mg/kg) were simultaneously injected intraperitoneal (IP). **(B)** Summary table of brain dialysate and serum pharmacokinetic measures. **(C)** Time course of estimated CSF (brain dialysate) concentration of TES2300 (black filled triangles, n=4) compared to TES2320 (orange filled triangles, n=4) in aCSF from 0-60 minutes post-IP injection. (B). **(D)** AUC of estimated CSF (brain dialysate) concentrations of TES2300 (black bars, n=4) compared to TES2320 (orange bars, n=4) in aCSF from 0-60 minutes post-IP injection. TES2320 showed a 1.43-fold increase over TES2300 in the CSF (brain dialysate). **(E)** AUC of estimated CSF (brain dialysate) concentrations of TES2320 (orange bars, n=4) normalized to TES2300 (black bars, n=4) in aCSF from 0-60 minutes post-IP injection. **(F)** Time course of plasma concentration of TES2300 (black filled triangles, n=4) compared to TES2320 (orange filled triangles, n=4) from 0-60 minutes post-IP injection. **(G)** AUC of plasma concentrations of TES2300 (black bars, n=4) compared to TES2320 (orange bars, n=4) from 0-60 minutes post-IP injection. TES2320 showed a 4.73-fold increase over TES2300 in the plasma. **(H)** AUC of plasma concentrations of TES2320 (orange bars, n=4) normalized to TES2300 (black bars, n=4) from 0-60 minutes post-IP injection. Data are presented as mean ± SEM. **p<0.01; ratio paired t-test.

#### Bioanalytical method for the detection of TES2300 and TES2320 peptide compounds in brain dialysate and plasma

Biological sample preparation: Plasma samples (20 µL) were mixed with 20 µL of an internal standard working solution containing 150 ng/mL (DADLE) in 10% acetic acid. To precipitate proteins in both plasma and dialysate, 120 µL of ice-cold methanol was added to the plasma mixture and the dialysate samples (20 µL, premixed with preservation solution). After vortexing for 30 seconds, samples were centrifuged at 8,000 x g for 10-minutes at 4°C. The supernatant was evaporated using a Savant Speedvac concentrator. Dialysate and plasma samples were reconstituted in 20 μL of diluent (5% methanol and 0.1% formic acid in nanopure water).

Liquid chromatography and mass spectrometry: Samples were injected into the Thermo Fisher Scientific UHPLC-MS/MS TSQ system. Separation was achieved on an ACQUITY UPLC HSS T3 C18 Column (100 Å, 1.8 µm, 2.1 × 100 mm) and VanGuard pre-column (HSS T3 1.8 μm, 2.1 × 5 mm) and a HyperSep Retain PEP trap column (2.1 × 10 mm) with the column temperature set at 40 °C. We used a water-acetonitrile gradient with a flow rate of 0.2 mL/min and a total run time of 25. The mobile phases used were A: 0.1% v/v formic acid in deionized water and B: 0.1% v/v formic acid in acetonitrile, with the following conditions: 0 to 1 min (2 %B diverted to waste), 1.0 to 1.2 min (5% B), 1.2 min to 6 min (15% B), 6 min to 11 min (30% B), 11 min to 12.5 min (95% B), 12.5 min to 18 min (95% B), 18 min to 18.5 min (5 %B) 18.5 min to 23.5 min (5% B), 23.5 min to 25 min (2 %B). Injection volumes of 10 µL for dialysate and 5 µL for the plasma sample were used. Detection was performed using a Thermo Fisher TSQ Fortis Plus mass spectrometer with an H-ESI ion source. The instrument was operated in positive-ion mode. The MRM mode was used for quantification, using target ions at m/z 477.88 [M + 6H]^6+^ → 589.25 (quantifier) and 552.50 (qualifier) for TES2300. Target ions at m/z 531.92 [M + 6H]^6+^ → 605.92 (quantifier), 716.75 (qualifier) for TES2320. Concentrations were calculated using the response factor-concentration plot. Pharmacokinetic parameters, including the area under the curve (AUC), maximum plasma concentration (Cmax), and time to reach Cmax (Tmax), were determined. CSF concentration is converted to an estimated concentration using the microdialysis probe recovery rate.

#### Repetitive mild traumatic brain injury model

At approximately PN56, we randomly divided mice into two groups: a repeated sham group and a repeated closed-head injury (CHI) group. The injuries were administered using an Impact One, Controlled Cortical Impact (CCI) Device, following previously established protocols(31–33). For the procedure, mice were first anesthetized with isoflurane (3.5% for induction, 2% for maintenance) and secured in a stereotaxic frame. The repetitive CHI model consisted of five separate concussive impacts delivered to the head at 24-hour intervals. Each impact was generated by an electromagnetically driven piston with a 4.0 m/s velocity, a 3 mm beveled, flat-tipped impactor, and a 1.0 mm depth with a 200 ms dwell time. The impact was precisely aimed at the bregma, which was visualized through the intact skin after hair removal. The sham group underwent identical procedures, including anesthesia and head fixation, but did not receive any impact. Throughout the procedure, we maintained the mice’s body temperature at 37°C using a warming pad. To minimize physiological stress, the total duration of isoflurane exposure and the entire surgical procedure was limited to no more than 5 minutes per session. Immediately after each sham or CHI procedure, we placed the mice in a clean cage on a warming pad and recorded the latency to self-right as an immediate post-injury assessment.

#### In situ hybridization (RNAscope)

We used the RNAscope in situ hybridization technique to quantify the expression of the gene encoding PACAP (*Adcyap1*) and PAC1R (*Adcyap1r1*) in the brains of male C57Bl/6J mice (n=7 sham, 8 mTBI). The following probes were obtained from Advanced Cell Diagnostics (Newark, CA): Mm-Adcyap1 (Cat# 405911) and Mm-Adcyap1r1 (Cat# 409561). We followed the standard manufacturer’s protocol for the RNAscope Fluorescent Multiplex Assay. The mice were first anesthetized via intraperitoneal injection of ketamine (85 mg/kg) and xylazine (10 mg/kg). They were then transcardially perfused with 1x phosphate-buffered saline (PBS) followed by 4% paraformaldehyde (PFA). The brains were dissected, post-fixed in 4% PFA for 24 hours, and then cryoprotected in a 20% sucrose solution for three days. The brains were then frozen on dry ice and stored at -80°C until sectioning. Using a cryostat, we cut coronal sections (20 µm thick) containing the LHb at two different anterior-posterior (AP) levels: approximately -1.20 mm and -1.50 mm caudal to bregma. The prepared sections were baked at 60°C, fixed in 4% PFA at 4°C, and then dehydrated using increasing concentrations of ethanol. This was followed by treatments with hydrogen peroxide, antigen retrieval, and Protease III. After each step, the sections were washed twice with Nanopure water for two minutes. We then performed RNA hybridization using antisense probes targeting the PACAP-encoding mRNA (*Adcyap1*) and PAC1R-encoding mRNA (*Adcyap1r1*) for two hours at 40°C. After a series of amplification steps (AMP 1-3), the sections were treated with the appropriate HRP conjugate and developed with Vivid dye (Vivid dye 650, 1:3000 dilution) and finally blocked. The sections were washed with Wash Buffer after each step. Lastly, the sections were cover slipped using Prolong Gold with DAPI. We acquired images using Zeiss AxioScan imaging. Quantification was performed using QuPath (v0.5.1), an established open-source software platform for digital pathology as used in our recent work(47). The workflow utilized:

1. **Nucleus Detection:** DAPI-stained nuclei were segmented using an intensity thresholding algorithm with standard nuclear parameters (cell radius, minimum/maximum nuclear area).
2. **Subcellular Spot/Puncta Detection:** Single-molecule mRNA transcripts (fluorescent puncta) were identified using QuPath’s built-in “Subcellular Spot Detection” plugin. Parameters were set with a fixed detection threshold above local background intensity, minimum spot distance, and expected spot size (0.5mm^2^).
3. **Metric Calculation:** The parameter “Average Puncta per Nucleus” calculates the total detected single-transcript spots within a defined anatomical region of interest (medial and lateral subregions of LHb, mLHb or lLHb) divided by the total number of positive DAPI nuclei in that same region.

#### Sucrose splash test

We conducted the sucrose splash test ∼20 days after the final mTBI or sham procedure to assess self-care grooming behavior as previously described(31, 33). This test is a useful measure of motivational deficits and anhedonia, as it quantifies an animal’s innate drive to maintain its cleanliness. Each mouse was placed individually into a clear cage and allowed to acclimate for 10 minutes. Following this baseline period, we gently removed the mouse and applied a 10% sucrose solution to its dorsal coat using two sprays from an atomizer. The sticky sucrose naturally prompts a vigorous self-grooming response. The mouse was immediately returned to the cage and monitored for an additional 5 minutes. All test sessions were video-recorded and analyzed by an experimenter blinded to the experimental conditions. We measured two primary parameters: Total duration of grooming: The total time (in seconds) the mouse spent actively grooming and Latency to groom: The time (in seconds) from the sucrose application until the first grooming bout began. Grooming was specifically defined as any active, continuous movement (lasting more than 3 seconds) involving the face, forelimbs, flank, or tail, such as touching, wiping, scrubbing, or licking.

#### Slice preparation

Electrophysiological experiments were conducted approximately four weeks post-mTBI, using separate cohorts of mice from those in the behavioral studies. Mice were deeply anesthetized with isoflurane and transcardially perfused with ice-cold, oxygenated artificial cerebrospinal fluid (ACSF). The ACSF solution contained (in mM): 126 NaCl, 21.4 NaHCO3, 2.5 KCl, 1.2 NaH2PO4, 2.4 CaCl2, 1.00 MgSO4, 11.1 glucose, and 0.4 ascorbic acid, and was continuously saturated with 95% O2-5% CO2. After perfusion, we rapidly removed the brain and kept it in ice-cold ACSF. Coronal slices (220 µm thick) containing the lateral habenula (LHb) were cut using a vibratome (Leica; Wetzler, Germany). These slices were then transferred to a recovery chamber and incubated in ACSF at 34°C for at least one hour before recordings. For patch-clamp recordings, individual slices were transferred to a submerged chamber and continuously perfused with ascorbic-acid free ACSF at 28-30°C.

#### Electrophysiology

We performed whole-cell and cell-attached patch-clamp recordings on LHb neurons in sagittal slices across the medial and lateral subregions of the LHb based on anatomical landmarks under infrared-differential interference contrast (IR-DIC) microscopy, rather than fluorescently tagged PAC1R-specific sub-populations. Because PAC1Rs are widely and densely expressed across the vast majority of LHb glutamatergic principal neurons (as confirmed by our RNAscope data in Figure 4 and Allen Brain Atlas expression profiles), random sampling of LHb neurons reliably captures the PAC1R-expressing population affected by mTBI. A MultiClamp 700B patch amplifier (Molecular Devices) and patch pipettes were used, with visualization via infrared-differential interference contrast microscopy. Data were acquired with a DigiData 1440A and pCLAMP 10 (Molecular Devices), filtered at 3 kHz, and digitized at 10 kHz. To assess spontaneous activity and excitability, we patched neurons with a potassium gluconate-based internal solution (130 mM K-gluconate, 15 mM KCl, 4 mM adenosine triphosphate (ATP)-Na^+^, 0.3 mM guanosine triphosphate (GTP)-Na^+^, 1 mM EGTA, and 5 mM HEPES, pH 7.28, 275-280 mOsm). Spontaneous action potential (AP) firing patterns were recorded at V=0 using cell-attached recordings in voltage-clamp mode for about 2 minutes. Spontaneous activity was categorized as: *silent:* no action potentials recorded during a 2-minute trace, *tonic:* continuous, regular or irregular AP firing without rhythmic grouping, and *bursting:* APs occurring in distinct clusters/groups separated by quiescent periods.

Neuronal excitability was evaluated by subjecting neurons to increasingly depolarizing current steps (10 pA to 100 pA, 5-second duration, 10 pA increments) while manually holding the membrane potential at - 65 to -70 mV. For each neuron, we counted the number of action potentials (APs) generated at each current step and averaged these values for generating I-V (current-voltage) plots. We also calculated the total APs per cell by summing the APs across all current steps. We also measured AP threshold, fast and medium after-hyperpolarization (fAHP and mAHP) amplitudes, AP half-width, and AP amplitude using Clampfit as previously described(31, 48, 49). Resting Membrane Potential (RMP) was measured in current-clamp mode immediately after achieving the whole-cell configuration. Input Resistance (Rin) was calculated by dividing the steady-state voltage response to a -50 pA current step by the current amplitude. Hyperpolarization-activated Cation Currents (Ih) was evoked by a series of 2-second voltage steps from - 50 mV to -120 mV. The amplitude of Ih was calculated as the difference between the peak and steady-state current. The associated sag potentials and rebound bursting were also measured in current-clamp mode. M-currents were recorded using a standard deactivation protocol where neurons were held at -60 mV, pre-pulsed to -20 mV, and then stepped to various potentials. The amplitude was the difference between the instantaneous and steady-state current trace. Throughout all experiments, we monitored both cell input resistance and series resistance. Any data where these values changed by more than 10% were excluded from the analysis.

#### Pharmacological Interventions

For all drug experiments, stock solutions of the PAC1R agonist TES2320 were prepared fresh daily in saline, which also served as the vehicle. Separate cohorts of sham and mTBI mice received intraperitoneal (i.p.) injections of either saline or TES2320 (10 mg/kg). Two distinct intervention strategies were employed: Early TES2320 intervention: Injections were administered within 30 minutes following each sham or mTBI injury. This early intervention was repeated five times with 24-hour intervals. Late TES2320 intervention: A single injection was given one-month post-sham or mTBI. Electrophysiology experiments and behavioral assessments were conducted four weeks post-mTBI.

#### Statistics

All data are presented as the mean ± Standard Error of the Mean (SEM), with statistical significance defined as p<0.05. All statistical computations were performed using GraphPad Prism 10. Specific statistical tests were applied as follows: A ratio paired t-test was used for BBB-penetration and pharmacokinetic analysis. Three-way ANOVA was used for the RNAscope data. Chi-square tests were employed to analyze the distribution of silent, tonic, or bursting LHb neurons during spontaneous activity across different experimental groups (sham vs. mTBI; vehicle vs. early TES2320 vs. late TES2320). Two-way ANOVAs were utilized for experiments involving depolarization-induced LHb excitability and I-V (current-voltage) plots. One-way ANOVA was used to assess differences in intrinsic passive and active membrane properties, as well as the latency and total grooming time data from the sucrose splash tests.

## Results

### Baseline Pharmacological Profile of TES2320

The glycopeptide TES2320 was engineered as a structurally constrained truncamer of PACAP_1-24_ bearing an O-linked β-lactoside serine amide at position 24 (Ser (*O*-Lactose)-NH_2_). The primary synthesis and foundational *in vitro* pharmacological screening of this compound were established earlier by Apostol et al.(27) and Smith et al.(29).

Utilizing label-free PWR spectroscopy, TES2320 was shown to maintain highly potent target engagement with human secretin-family GPCRs. TES2320 exhibits picomolar binding at the human PAC1 receptor (K_d_ = 4.50 pM). These receptors were extracted from CHO cells and embedded in egg phosphatidyl choline (PC) membranes. Additionally, the compound exhibits macro-affinity partitioning into receptor-free lipid bilayers (EggPC K_d_ = 1.35 nM), a parameter influenced by the amphipathic properties introduced via C-terminal glycosylation.

#### Effects of PACAP glycosylation on blood-brain barrier penetration and pharmacokinetic measures

To demonstrate that the lactoside-bearing drug TES2320 was being delivered to the brain and determine pharmacokinetic changes in the periphery, it was administered to mice simultaneously with its unglycosylated analogue TES2300 *via* a catheterized carotid artery. Blood samples were removed every five minutes from the catheter, and microdialysis from the brain (Figure 2A-D) was time-locked (5 minute bins) to plasma (Figure 2 A, E-G) collected from C57Bl/6J mice post IP injection. In dialysate, the mean AUC of TES2300 was 703 ± 338 ng•min/mL, the Cmax was 64 ± 21 ng/mL, and the Tmax was 8 minutes post-injection. For the dialysate, the mean AUC of TES2320 was 1071 ± 574 ng•min/mL, with a mean Cmax of 74 ± 22 ng/mL. Tmax for the glycosylated PACAP was also 8 minutes. TES2320 exposure was observed in the plasma, with a mean AUC of 6089 ± 1370 ng·min/mL and a Cmax of 313 ± 62 ng/mL, and the Tmax was ∼10 minutes post-injection (n = 4) which was significantly larger than the unglycosylated analogue; TES2300 dose exposure in the plasma was observed with a mean AUC of 1370 ± 418 ng·min/mL, a Cmax of 106 ± 25 ng/mL, and a Tmax of ∼ 4 minutes (n = 4). The AUC C_brain_/C_plasma_ of -0.176 indicates good brain penetration of TES2320, and is in the range of other glycoside drug candidates, such as PNA5, an angiotensin 1-7 glucoside developed for the treatment of cognitive decline (AUC C_brain_/C_plasma_: 0.255) (50). To account for sample-to-sample baseline differences, we utilized a ratio paired t-test comparing TES2300 and TES2320 within the same time-locked biological runs, demonstrating statistically robust increases in exposure (Figure 2D–H). By co-injecting the drugs simultaneously in each animal we not only reduce the number of mice used, but also eliminate sources of error due to animal-to-animal variation, variation in timing, and injection site variation. Because of the high affinity for the membrane displayed by both TES2300 and TES2320 (K_d_’s of 0.114 nM and 14.1 nM, respectively), microdialysis can only deliver an estimate of the true amount of active peptide in the brain since the vast majority of these species adsorb to the cellular membranes in the brain, which are vast relative to the brain volume.

#### Effects of mTBI on PACAP and PAC1R mRNA expression in the LHb

Previous research has demonstrated that activating PACAP-expressing neurons in the LHb—identified by the *Adcyap1* gene and a glutamatergic marker (VGLUT2(11))-produces unexpected behavioral outcomes associated with reward, reduced anxiety, and diminished fear(16). Given that our mTBI model induces LHb hyperactivity and motivational deficits, we hypothesized that mTBI might cause a PACAP deficiency by reducing PACAP expression specifically in these neurons which may also alter PAC1R expression within the LHb. To test this, we used multiplex fluorescent in situ hybridization (RNAscope) to quantify PACAP and PAC1R mRNA expression in the medial and lateral subregions of the LHb at two different anterior-posterior (AP) levels. Our results confirmed this hypothesis: irrespective of subregion or AP level, mTBI significantly decreased PACAP mRNA levels in the LHb (Figure 3, sham vs. mTBI, F (1, 52) = 7.117, *p<0.05, three-way ANOVA) while increasing PAC1R mRNA levels (Figure 4, sham vs. mTBI, F (1, 50) = 5.869, *p<0.05, three-way ANOVA). Overall, our findings reveal a significant up-regulation of PAC1R expression in the LHb following mTBI. This increase in the receptor could represent a compensatory, homeostatic response to the persistent decrease in PACAP mRNA that we observed after mTBI.

**Figure 3.**
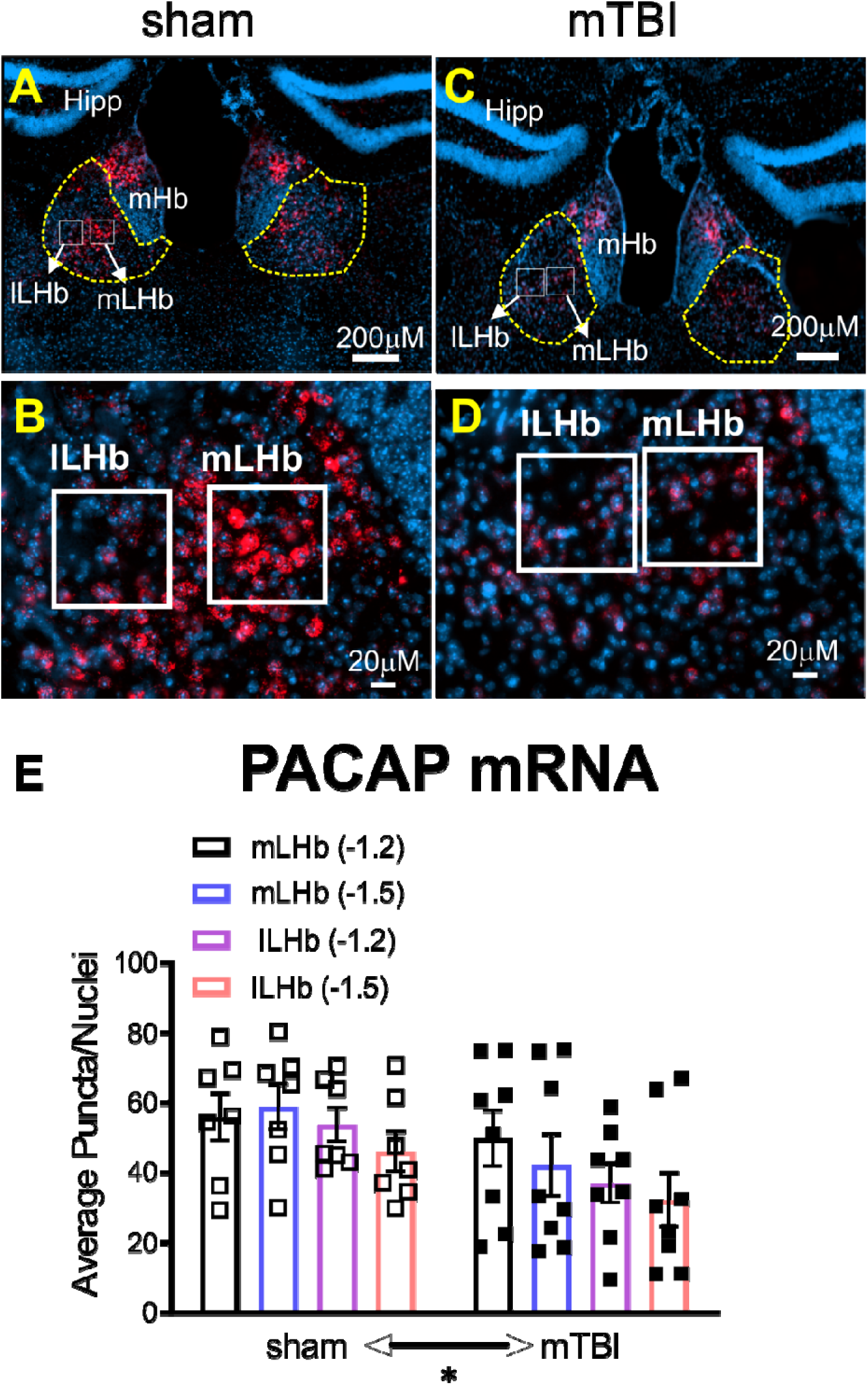
mTBI leads to a significant reduction in PACAP expression in the mouse LHb. This figure uses RNAscope technology to show the effect of mTBI on the expression of PACAP mRNA in the mouse brain. (**A-D**) Representative images from sham and mTBI mice (A-P level: -1.2 relative to bregma). Panels **A** and **C** are low-magnification overviews of the habenula, showing the medial (mHb) and lateral (LHb) regions. Panels **B** and **D** are high-magnification images showing individual PACAP mRNA puncta (red) localized within the medial LHb (mLHb) and lateral LHb (lLHb) subregions. The cell nuclei are stained with DAPI (blue). Scale bars for low and high magnification are 200µm and 20µm, respectively. (**E**) Quantitative analysis shows a significant decrease in PACAP mRNA signal (puncta per nucleus) in the LHb (A-P levels: -1.2, -1.5 relative to bregma) of mTBI mice compared to sham controls. Data are presented as the mean ± SEM from two different AP levels. *p<0.05; three-way-ANOVA.

**Figure 4.**
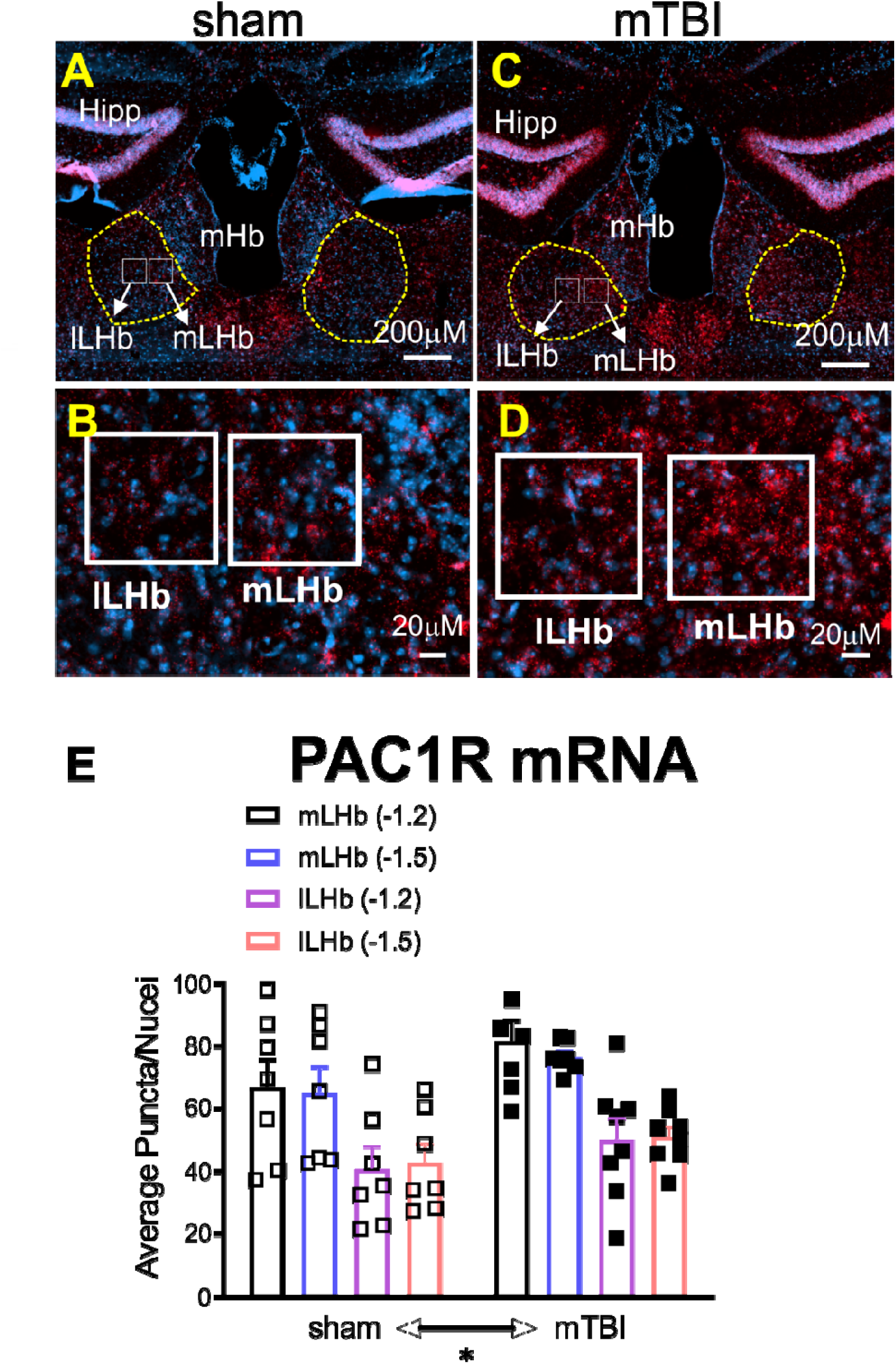
mTBI leads to a significant increase in PAC1R expression in the mouse LHb. This figure uses RNAscope technology to show the effect of mTBI on the expression of PAC1R mRNA in the mouse brain. (**A-D**) Representative images from sham and mTBI mice (A-P level: -1.2 relative to bregma). Panels **A** and **C** are low-magnification overviews of the habenula, showing the medial (mHb) and lateral (LHb) regions. Panels **B** and **D** are high-magnification images showing individual PAC1R mRNA puncta (red) localized within the medial LHb (mLHb) and lateral LHb (lLHb) subregions. The cell nuclei are stained with DAPI (blue). Scale bars for low and high magnification are 200µm and 20µm, respectively. (**E**) Quantitative analysis shows a significant increase in PAC1R mRNA signal (puncta per nucleus) in the LHb (A-P levels: -1.2, -1.5 relative to bregma) of mTBI mice compared to sham controls. Data are presented as the mean ± SEM from two different AP levels. *p<0.05; three-way-ANOVA.

#### Effects of early and late TES2320 intervention on LHb spontaneous activity and neuronal excitability following mTBI

To determine if early or late TES2320 intervention could prevent or reverse mTBI-induced LHb hyperactivity, we examined several neurophysiological parameters including spontaneous activity (Figure 5A), neuronal excitability (Figure 5B-C), intrinsic membrane properties (Figure 6), hyperpolarization-induced rebound bursting and two membrane ionic currents (Ih currents and M-currents) (Figure 7), in LHb slices from 4 separate groups of male mice : sham or mTBI mice receiving vehicle as well as mTBI mice receiving either early or late TES2320 treatment at ∼4 weeks post-injury. Consistent with our previous findings(31–33), mTBI led to a persistent increase in spontaneous LHb tonic activity in LHb neurons from vehicle-treated mTBI mice compared to those from vehicle-treated sham mice. Both early and late PAC1R agonist treatments in mTBI mice successfully prevented and reversed this hyperactivity, respectively (Figure 5A, sham+vehicle vs. mTBI+vehicle, **p<0.01, mTBI+vehicle vs. mTBI+early TES2320, **p<0.01, mTBI+vehicle vs. mTBI+late TES2320, *p<0.05, Chi squared test). Next, we performed neuronal excitability recordings where we evaluated action potential (AP) generation in response to depolarization. As previously observed(31), LHb neurons from vehicle-treated mTBI mice fired more APs in response to depolarization without any changes in intrinsic properties compared to vehicle-treated sham mice. Notably, both early and late TES2320 treatments in mTBI mice normalized this hyperexcitability to sham levels (Figure 5B, I-V plots: sham+vehicle vs. mTBI+vehicle, F (1, 560) = 49.72, ****P<0.0001, mTBI+vehicle vs. mTBI+early TES2320, F (1, 1077-) = 73.14, ****P<0.0001, mTBI+vehicle vs. mTBI+late TES2320, F (1, 508) = 46.72, ****P<0.0001, 2-way ANOVA; Figure 5C, Sum of APs: sham+vehicle vs. mTBI+vehicle, *p<0.05, mTBI+vehicle vs. mTBI+early TES2320, **p<0.01, mTBI+vehicle vs. mTBI+late TES2320, *p<0.05, one-way ANOVA). The only exception in intrinsic properties was the fAHP, which was significantly diminished in the mTBI+late TES2320 group compared to the mTBI+vehicle group (Figure 6, mTBI+vehicle vs. mTBI+late TES2320, *p<0.05, one-way ANOVA).

**Figure 5:**
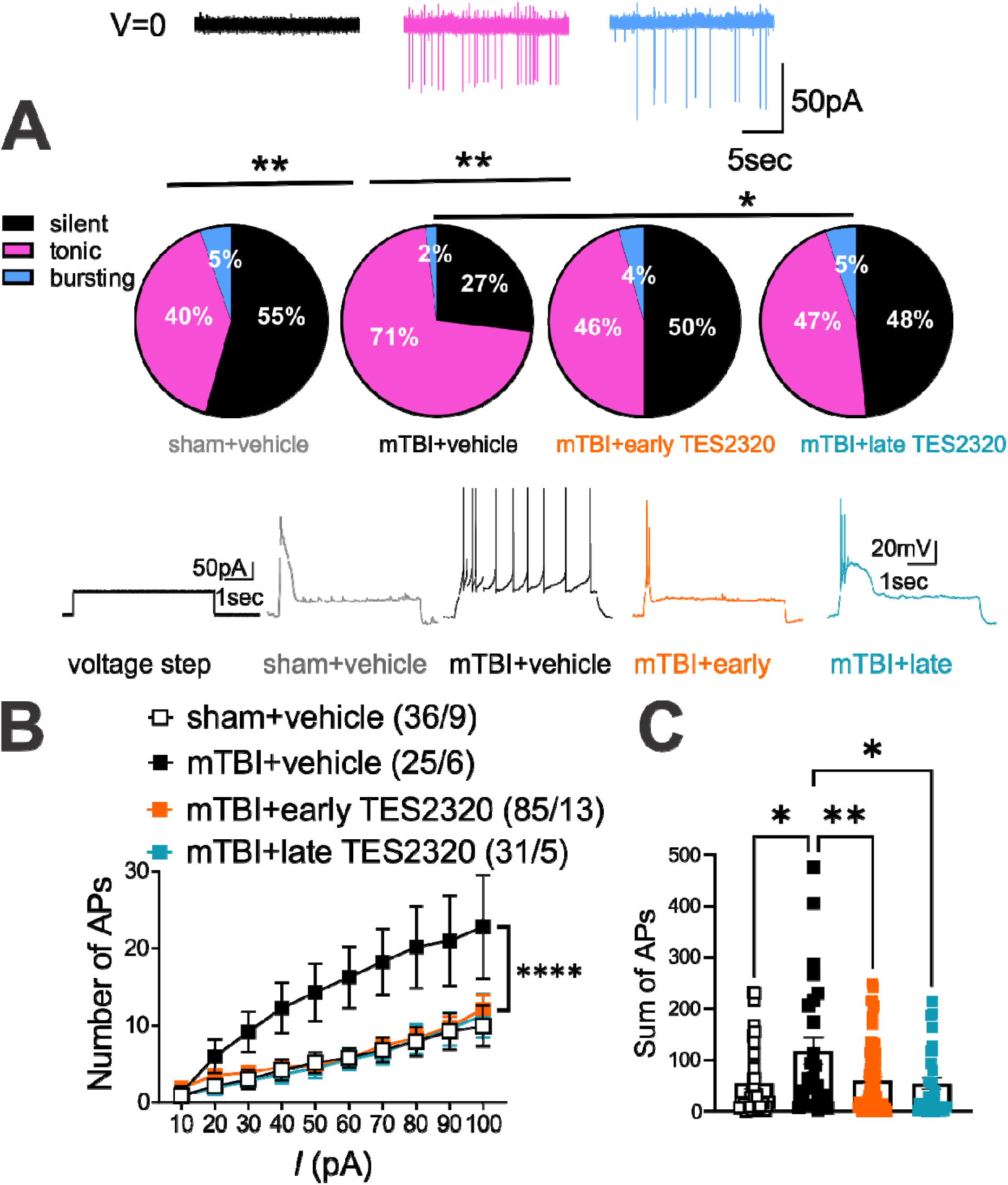
TES2320 treatment normalizes the increased spontaneous activity and excitability of LHb neurons in male mTBI mice. (**A**) Spontaneous neuronal firing patterns of LHb neurons. Representative voltage-clamp traces and corresponding pie charts illustrate the percentage of cells classified as silent (black), tonic (magenta), or bursting (blue) from sham+vehicle (n=52 cells/9 mice), mTBI+vehicle (n=52 cells/6 mice), mTBI+early TES2320 (n=138 cells/13 mice), and mTBI+late TES2320 (n=58 cells/5 mice) mice. Recordings were performed in cell-attached configuration at a holding potential of 0 mV. (**B-C**) Neuronal excitability of LHb neurons. Action potential (AP) generation in response to depolarizing current steps and a summary graph of the total number of APs per cell are shown. Representative traces (in response to a 50pA current step) are from sham+vehicle (grey trace, black open squares, n=36 cells/9 mice), mTBI+vehicle (black trace, black filled squares, n=25 cells/6 mice), mTBI+early TES2320 (orange trace, orange filled squares, n=85 cells/13 mice), and mTBI+late TES2320 (turquoise trace, turquoise filled squares, n=31 cells/5 mice) male mice. Recordings were made under conditions of intact synaptic transmission. Data are presented as mean ± SEM. *p<0.05 and ****p<0.0001; Chi-square test (for A) and one-way and two-way ANOVA (for B-C).

**Figure 6:**
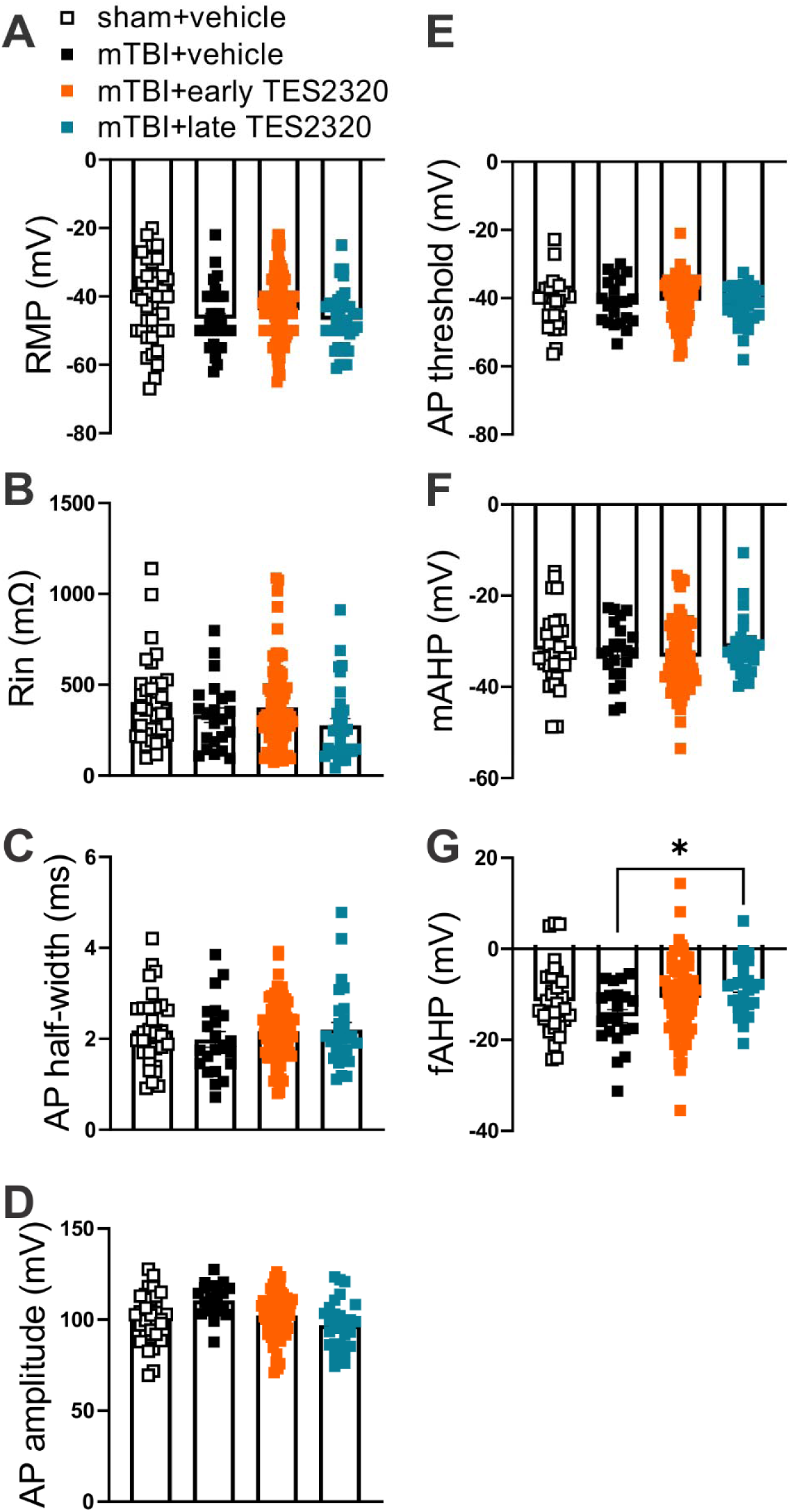
Late TES2320 treatment decreases the fast afterhyperpolarization (fAHP) amplitude in mTBI mice. (**A-G**) Electrophysiological properties of LHb neurons. Summary graphs of resting membrane potential (RMP), input resistance (Rin), AP half-width, AP amplitude, AP threshold, medium afterhyperpolarization (mAHP), and fast afterhyperpolarization (fAHP). All parameters were calculated from depolarization-induced AP recordings, as depicted in Figure 5. Data are from sham+vehicle (black open squares, n=36 cells/9 mice), mTBI+vehicle (black filled squares, n=25 cells/6 mice), mTBI+early TES2320 (orange filled squares, n=85 cells/13 mice), and mTBI+late TES2320 (turquoise filled squares, n=31 cells/5 mice) male mice. Data are presented as mean ± SEM.*p<0.05; one-way ANOVA.

**Figure 7:**
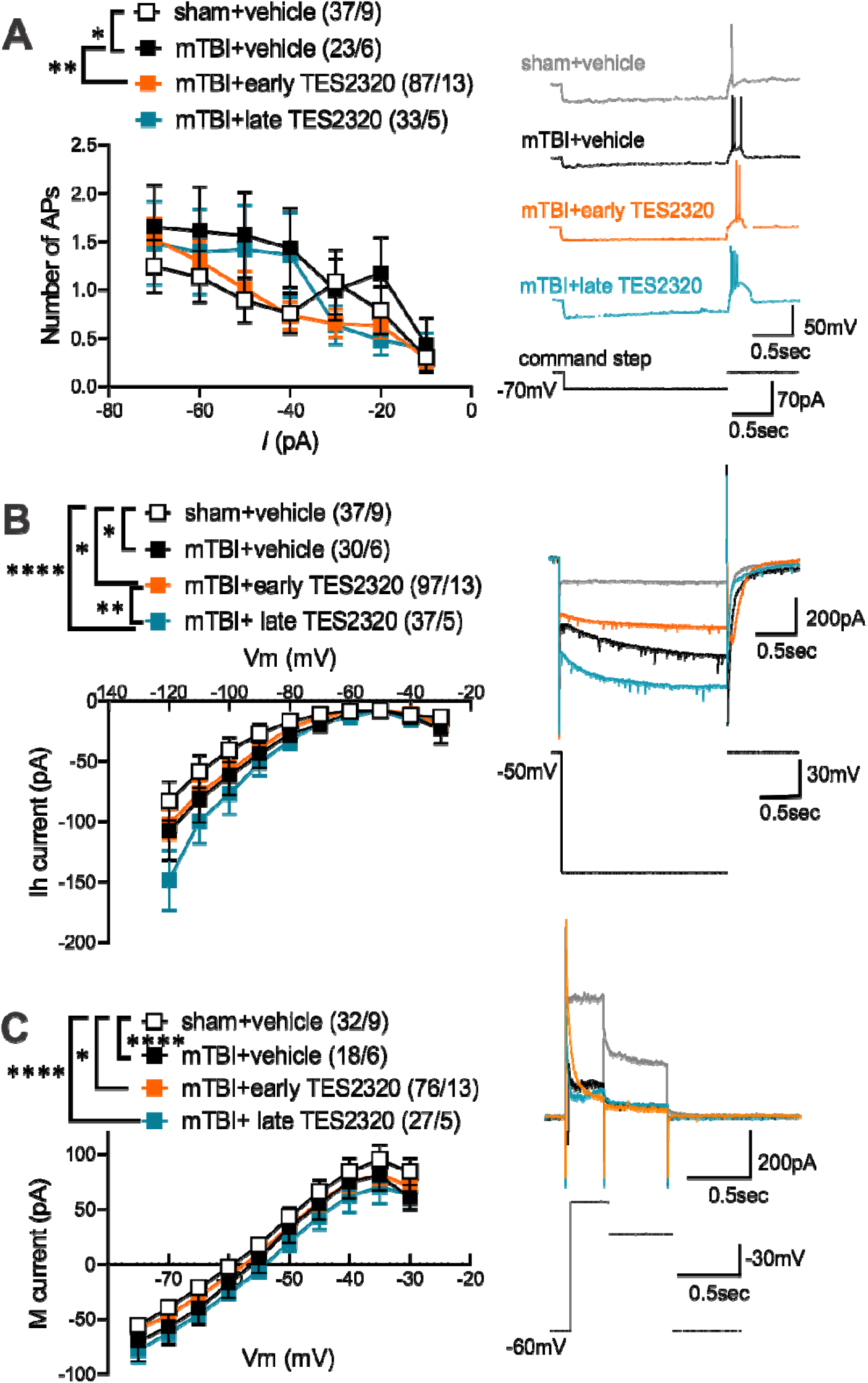
Early TES2320 treatment normalizes mTBI-induced increases in hyperpolarization induced rebound bursts, while late TES2320 further enhances Ih currents in LHb neurons. (**A**) Hyperpolarization-induced rebound bursts. A summary graph of the average number of APs in rebound bursts triggered in response to hyperpolarizing current steps in current-clamp recordings in LHb neurons is shown. Representative traces of the rebound bursts in response to a -70pA hyperpolarizing current injection are from LHb neurons of sham+vehicle (grey trace), mTBI+vehicle (black trace), mTBI+early TES2320 (orange trace), and mTBI+late TES2320 (turquoise trace) male mice. Data were recorded using current-clamp. (**B**) Ih currents. The I-V relationship of hyperpolarization-activated inward currents (Ih) and representative voltage-clamp traces at -120mV voltage step are shown with similar color coding of traces as in (A). (**C**) M-currents. The I-V relationship of M-currents and representative voltage-clamp traces at -30mV voltage step are shown with similar color coding of traces as in (A). Data in all panels are from sham+vehicle (black open squares, n=37 cells/9 mice in A, n=37 cells/9 mice in B, n=32 cells/9 mice in C), mTBI+vehicle (black filled squares, n=23 cells/6 mice in A, n=30 cells/6 mice in B, n=18 cells/6 mice in C), mTBI+early TES2320 (orange filled squares, n=87 cells/13 mice in A, n=97 cells/13 mice in B, n=76 cells/13 mice in C), and mTBI+late TES2320 (turquoise filled squares, n=33 cells/5 mice in A, n=37 cells/5 mice in B, n=27 cells/5 mice in C) male mice. Data are presented as mean ± SEM. *p<0.05, **p<0.01, and ****p<0.0001; two-way ANOVA.

#### Effects of early and late TES2320 intervention on hyperpolarization-induced rebound bursting and ion currents in LHb neurons following mTBI

Our previous work(31) also showed that LHb neurons from mTBI mice displayed less spontaneous bursting but more rebound bursts in response to hyperpolarization compared to those from sham mice(31). We observed the same effect in our vehicle-treated groups. Interestingly, only early TES2320 intervention was able to normalize this mTBI-induced increase in rebound bursting in LHb neurons from early TES2320-treated mTBI mice compared to those from vehicle-treated mTBI mice. Late TES2320 treatment had no effect on this measure (Figure 7A, sham+vehicle vs. mTBI+vehicle, F (6, 406) = 2.616, *p<0.05, mTBI+vehicle vs. mTBI+early TES2320, F (1, 756) = 7.277, **p<0.01). We then examined two key ion currents, Ih currents and M-currents, both of which play a significant role in regulating LHb activity and bursting(48, 51–56). Ih was calculated as the difference between the peak initial inward current and the steady-state current at the end of a 2-second hyperpolarizing step (from -50mV to -120 mV). M-current amplitude was measured during a deactivation protocol as the difference between the instantaneous peak and steady-state relaxed current trace during voltage steps from -20 mV. As previously reported(31), mTBI enhanced Ih currents in LHb neurons as evident from comparison of Ih currents of LHb neurons in vehicle-treated mTBI mice versus those from sham mice receiving vehicle. Surprisingly, early TES2320 was insufficient to prevent this Ih enhancement in LHb neurons following mTBI. Furthermore, late TES2320 treatment exaggerated this effect, resulting in even larger Ih currents in LHb neurons from late TES2320-treated mTBI mice than those from the sham and early TES2320 groups (Figure 7B, sham+vehicle vs. mTBI+vehicle : F (1, 660) = 6.21, **p<0.01, sham+vehicle vs. mTBI+early-TES2320, F (1, 1330) = 4.569, *p<0.05, sham+vehicle vs. mTBI+late TES2320+, F (1, 730) = 17.98, ****p<0.0001, mTBI+early TES2320 vs. mTBI+late TES2320, F (1, 1320) = 8.595, *p<0.05, 2-way ANOVA). Consistent with earlier findings(31), mTBI significantly reduced M-currents in LHb neurons from vehicle-treated mTBI mice compared to those from vehicle-treated sham mice. Neither early nor late TES2320 intervention in mTBI mice normalized this reduction (Figure 7C, sham+vehicle vs. mTBI+vehicle : F (3, 1489) = 7.216,****p<0.0001, sham+vehicle vs. mTBI+early-TES2320, F (1, 1059) = 4.386, *p<0.05, sham+vehicle vs. mTBI+late TES2320+, F (1, 570) = 25.33, ****p<0.0001, 2-way ANOVA).

#### Effects of early and late TES2320 interventions on grooming behaviors following mTBI

Consistent with our previous findings(33), vehicle-treated mTBI mice showed a significantly increased latency to groom during the sucrose splash test compared to sham mice, indicating a motivational deficit. However, their total grooming time was unchanged. Only early TES2320 treatment in mTBI mice was effective in preventing the prolonged grooming latency caused by mTBI. In contrast, late TES2320-treated mTBI mice continued to exhibit increased grooming latency compared to both the sham and early TES2320 groups. Furthermore, late TES2320 treatment in mTBI mice significantly decreased the total grooming time compared to vehicle-treated sham mice and mTBI mice that received early TES2320 treatment (Figure 8, latency: sham+vehicle vs. mTBI+vehicle, *p<0.05, mTBI+vehicle vs. mTBI+early-TES2320, *p<0.05, sham+vehicle or mTBI+early TES2320 vs. mTBI+late TES2320, **p<0.01; total grooming: sham+vehicle vs. mTBI+late TES2320, *p<0.05, mTBI+early TES2320 vs. mTBI+late TES2320, **p<0.01, one-way ANOVA).

**Figure 8:**
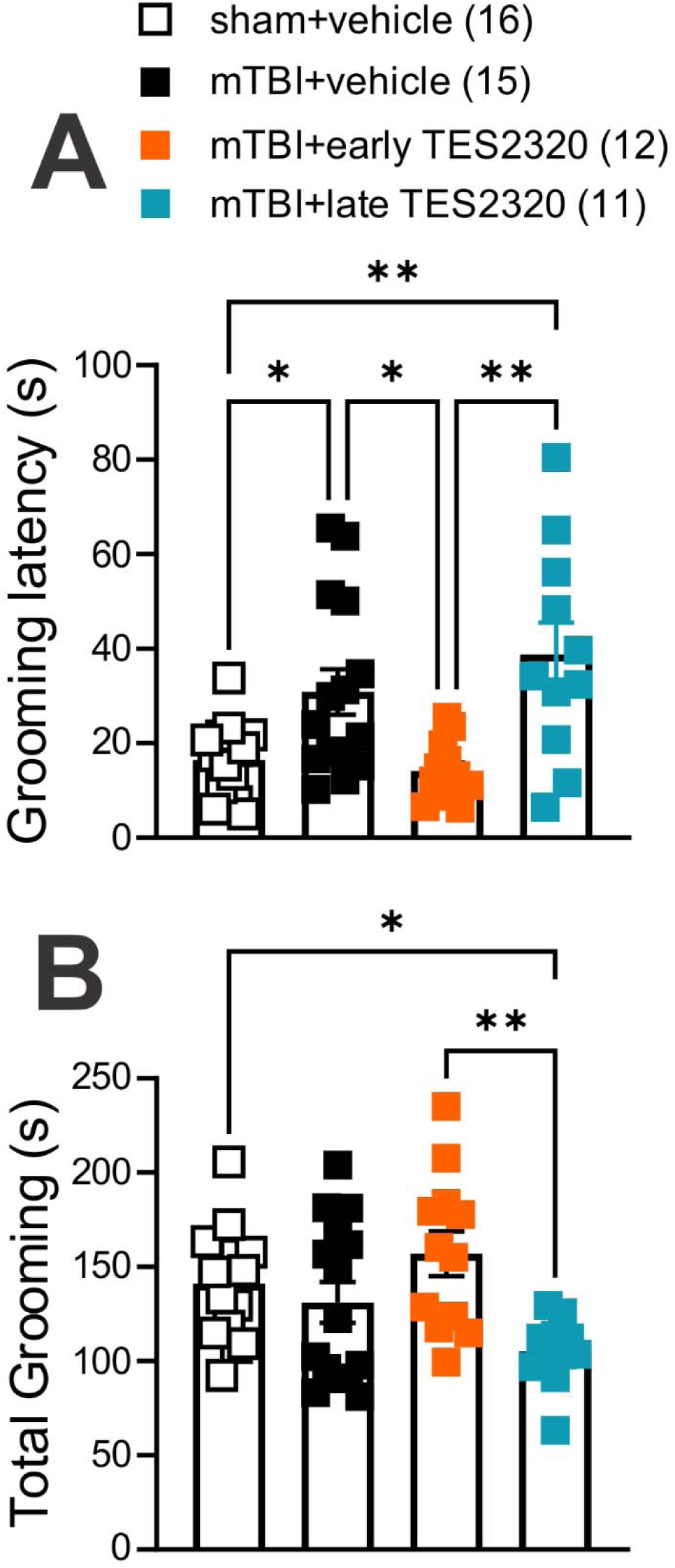
Early TES2320 treatment reverses mTBI-induced motivational deficits in self-grooming behavior. (**A**) Latency to groom. A graph showing the time to initiate grooming behavior in the sucrose splash test. Early TES2320 treatment normalized the increased latency to groom seen in mTBI mice. (**B**) Total grooming time. A graph showing the total time spent grooming during the sucrose splash test. While early TES2320 treatment had no effect, late TES2320 treatment significantly decreased total grooming time in mTBI mice compared to both sham and early TES2320-treated mice. All data are from male mice: sham+vehicle (black open squares, n=16) mTBI+vehicle (black filled squares, n=15), mTBI+early TES2320 (orange filled squares, n=12) and mTBI+late TES2320 (turquoise filled squares, n=11). Data are presented as mean ± SEM. *p<0.05, **p<0.01; one-way ANOVA.

## Discussion

This study addresses a critical gap in mTBI research by exploring the long-term efficacy of a novel PACAP glycopeptide analogue with high affinity for PAC1R, TES2320, in a closed-head injury mouse model. Our investigation began by confirming that mTBI leads to a significant reduction in PACAP mRNA expression in the LHb, a region we have identified as critical for the motivational deficits observed after injury(33). The LHb is a subcortical anti-reward region whose activity is altered by our repetitive mTBI model, leading to hyperactivity, enhanced rebound bursting, and a lack of motivation in self-care, disrupted social behavior, and aberrant threat responses(31–33). These symptoms are core to many psychiatric conditions, including depression, anxiety, and PTSD.

Our previous studies also showed that mTBI promotes the dysregulation of key neuromodulatory stress systems, including the corticotropin-releasing factor/CRFR1 (CRF/CRFR1) and dynorphin/kappa opioid receptor (DYN/KOR) systems, which both are known to interact with the PACAP system(12, 13, 57–61). PACAP and PAC1Rs are highly expressed in stress-related neural circuits(8, 10–12), including within intrinsic LHb PACAP-expressing neurons (11, 16). Notaby, the role of PACAP in mood is complex and brain region-specific; for example, while systemic PACAP can decrease motivation and social behaviors(62), optogenetic activation of PACAP-expressing neurons in the hippocampal dentate gyrus triggers antidepressant responses(63). More importantly, emerging evidence suggests that intrinsic LHb PACAP-expressing neurons exert an unexpected anxiolytic and rewarding influence (16), challenging the traditional view of the LHb as a purely aversive circuit.

A likely mechanism underlying this counterintuitive phenotype is that these glutamate- and PACAP-co-releasing neurons possess distinct local connectivity that starkly contrasts with classical LHb aversion pathways. Thus, the physiological outcome of glutamate and PACAP co-release appears to be circuit dependent. For instance, local PACAPergic neurons may provide excitatory drive onto LHb interneurons; their activation would subsequently suppress downstream LHb glutamatergic projection neurons, yielding the anxiolytic and antidepressant effects reported by Levinstein and colleagues(16). Given the presence of such local PACAPergic microcircuits and high postsynaptic PAC1R expression within the LHb (11, 16), supplemented by dense extrinsic PACAPergic inputs to the LHb from hypothalamic stress centers (e.g., PVN, lateral hypothalamus) and anterior insular/mPFC regions, the mTBI-induced reductions in LHb PACAP mRNA likely reflect a simultaneous disruption of both local microcircuits and long-range projections. This chronic signaling deficit may trigger a compensatory upregulation of postsynaptic PAC1Rs within the LHb.

While systemic TES2320 administration broadly engages central PAC1Rs, two lines of evidence establish the LHb as a primary functional substrate for its therapeutic actions: our prior finding that chemogenetic silencing of LHb glutamatergic neurons rescues mTBI-induced grooming deficits in the sucrose splash test (33), and our current electrophysiological data showing that TES2320 treatment directly dampens LHb hyperexcitability *ex vivo*. Mechanistically, several passive intrinsic membrane properties, as well as the mTBI-induced enhancement of I*_h_* currents, remained largely unchanged following TES2320 treatment. However, because these recordings were conducted in the presence of intact synaptic transmission, ongoing local network activity may mask subtle postsynaptic baseline alterations. Furthermore, postsynaptic PAC1R activation is known to recruit intracellular cascades (G*_αs_*/G*_αq_*)(64) that can drive the synthesis and retrograde release of messengers such as nitric oxide(65, 66), which selectively tune presynaptic neurotransmitter release without altering somatic baseline conductances.

Importantly, despite the persistence of elevated I*_h_* currents, early treatment with TES2320 effectively normalized mTBI-induced LHb rebound bursting. Because I*_h_* -mediated depolarizing sag interacts with low-threshold T-type calcium channels (Ca_V_3) and postsynaptic NMDAR activation to facilitate rebound burst generation in LHb neurons(53), this selective rescue suggests that early PACAP receptor activation acts downstream of I*_h_*. Specifically, TES2320 may normalize hyperexcitability by targeting T-type Ca^2+^ channel gating or NMDAR-dependent burst-generating cascades, rather than acting through non-specific suppression of all intrinsic ionic conductances. Together, these findings indicate that TES2320’s therapeutic efficacy involves targeted modulation of specific burst-promoting intrinsic mechanisms and synaptic inputs within vulnerable LHb circuits

Based on this biophysical framework, we evaluated the long-term therapeutic potential and timing-dependence of TES2320. We found that both early and late interventions effectively prevented and reversed the tonic LHb hyperactivity and hyperexcitability caused by mTBI. However, the timing of treatment was a determining factor for complete functional and behavioral recovery. Only early intervention successfully prevented the enhanced rebound bursting in LHb neurons and normalized grooming latencies, suggesting that mTBI-induced motivational deficits in self-care are responsive to a timely PAC1R agonist intervention. In contrast, late TES2320 intervention failed to reverse grooming latencies and acutely decreased overall grooming behavior. These negative outcomes are likely a result of the acute, single-dose nature of the late intervention, which was evaluated within a 24-hour timeframe, a period known for PACAP’s persistent side effects on mood and motivation lasting up to 7 days(62, 67). This contrasts sharply with the early intervention protocol, which involved repeated doses but was evaluated a month post-injury, well beyond the window for any acute side effects. This difference in experimental design, specifically the timing and duration of the intervention, warrants further investigation into the dose-dependent effects of TES2320 and their impact on long-term behavioral outcomes. Both native PACAP and its engineered analogues exhibit high affinity for PAC1Rs, which we have confirmed are highly expressed within the LHb. It is therefore highly probable that the neurophysiological actions of TES2320 on LHb neurons are mediated through PAC1R activation. However, unlike native PACAP, TES2320 also displays picomolar functional potency at VPAC1 receptors (29). While previous literature and the *Allen Mouse Brain Reference Atlas* show that LHb VPAC1 expression is relatively low compared to PAC1R (68, 69), VPAC1 signaling may still contribute to the therapeutic benefits or behavioral side effects of TES2320.

## Concluding Remarks and Future Directions

The present study demonstrates that TES2320, a novel PAC1R agonist, offers a promising therapeutic strategy for mitigating the long-term neurophysiological and behavioral deficits following mTBI. Our findings underscore that the timing of this intervention is critical, with early administration successfully preventing rebound bursting and restoring motivational behaviors. These results suggest that TES2320, and similar glycosylated PACAP analogues with additional truncation, possess powerful antidepressant-like effects in addition to their established neuroprotective and anti-inflammatory properties in animal models of TBI. Overall, our work validates glycosylation as a key method for developing viable peptide drug candidates for the treatment of mTBI-related disorders.

Several limitations of this study warrant consideration. First, quantifying brain concentrations of TES2320 is technically challenging due to the strongly amphipathic, surfactant-like nature of the PACAP helix. Consequently, microdialysis measures only the free drug in the aqueous extracellular compartment, largely underrepresenting peptide adsorbed to cellular membranes or dialysis tubing. Nevertheless, these measurements confirm that substantial levels of TES2320 persist in serum and cross the blood–brain barrier (BBB) following peripheral administration.

Second, while the sucrose splash test provides a benchmark, causally validated measure of LHb-dependent self-care motivation in our repetitive mTBI model(33), we acknowledge that evaluating a broader battery of affective and cognitive behaviors will be important to fully map the scope of TES2320’s therapeutic rescue across all behavioral domains in future studies.. Furthermore, the selection of the 10mg/kg (i.p.) dose in the current study was informed by extensive prior pharmacokinetic and efficacy studies with related PACAP glycopeptide analogues (e.g., 2LS80Mel and PNA5) in preclinical models of stroke and traumatic brain injury (9, 50), which confirmed that this dose reliably yields neuroprotective central concentrations without systemic toxicity. Nevertheless, comprehensive dose-response evaluations paired with an expanded behavioral battery will be essential to fully establish the optimal therapeutic window and clinical utility of TES2320.

Finally, this study evaluated TES2320 exclusively in male mice. Human studies have suggested that dysregulation of the PACAP–PAC1R pathway contributes to abnormal stress responses and mediates part of the sexual dimorphism observed in stress-related disorders like PTSD, to which females are more predisposed(25, 70–72). Although our mTBI model generally produces similar physiological and behavioral outcomes in young adult male and female mice, we have detected notable sex differences. For example, female mice exhibit compensatory upregulation of M-currents in LHb neurons, and while both sexes show a shift toward passive, action-locking defensive behaviors, females have prolonged latencies to escape, suggesting a greater predisposition to PTSD-related phenotypes(31). Considering the high expression of estrogen receptors in the LHb and the ability of estradiol to suppress LHb neuronal activity(73–75), it is plausible that estrogen-mediated alterations in PACAP signaling could contribute to sexual dimorphism in LHb circuits and their responsivity to the antidepressant effects of PACAP analogues, specifically PAC1R agonists(76). Our future work will therefore include female mice and delve into circuit-behavioral studies to delineate how mTBI alters PACAP-expressing LHb circuits and their interaction with the CRF and DYN/KOR neuromodulatory systems. Together, our current findings indicate that TES2320’s efficacy stems, at least in part, from restoring physiological PACAP signaling within vulnerable LHb circuits.

## Acknowledgments

The opinions and assertions contained herein are the private opinions of the authors and are not to be construed as official or reflecting the views of the Uniformed Services University of the Health Sciences or the Department of War or the Government of the United States. Behavioral testing and analysis were performed in the Preclinical Behavior and Modeling Core at the Uniformed Services University. The authors acknowledge the use of Biorender for Figure 2 (https://BioRender.com/i4s4zn8) and Gemini as a tool for editing the manuscript, with the final content and original draft remaining the sole work of the authors.

## Authorship contribution

The study concept and design were developed by F.N. and R.P. The experimental design for the mild traumatic brain injury (mTBI) was specifically guided by R.A. The PAC1R agonist was designed and synthesized by T.S., L.S., F.A.A. and R.P., E.T. and W.F. carried out the sham and mTBI procedures. Data acquisition was performed by E.T., S.G., W.F., C.O., M.C., M.J.B., and F.N., with E.T., S.G., O.R., B.C., R.A., T.F., M.L.H., C.O., M.J.B., and F.N. assisting in data analysis and interpretation. The initial manuscript draft was prepared by E.T., R.P., and F.N. All authors critically reviewed and approved the final manuscript for submission. The authors extend their gratitude to Dr. Yeonho Kim, Dr. Amanda Fu, and Laura Tucker at the Uniformed Services University (USU) Preclinical Modeling and Behavior Core for their instrumental support in the implementation of the sham and mTBI procedures and behavioral studies.

## Conflict of Interest statement

Polt, M. L. Heien and T. Falk have a financial interest in *Teleport Pharmaceuticals,* which expects to license the PACAP glycosides from Tech Launch Arizona, the patent holder. The other authors have no competing interests to declare.

## Funding statement

This work was supported by the Congressionally Directed Medical Research Programs (CDMRP) under Grant #HT9425-23-2-0003 to F.N. and R.P. The funding agency did not contribute to writing this article or deciding to submit it.

## Transparency, Rigor, and Reproducibility Statement

For all experiments, animals were randomly assigned to groups, and all data acquisition and analysis were performed by investigators blinded to the group assignments (sham/mTBI and vehicle/drug). This ensured the scientific rigor and reproducibility of the findings. The microdialysis and pharmacokinetic study’s sample size (n = 4 mice per group) and *in vivo* TBI studies (n = 5–16 mice per group) were determined to detect a significant effect of injury, drug, and their interactions, with statistical significance set at p<0.05. Statistical analyses were performed using a ratio paired t-test, Chi-square tests and one-, two-, or three-way ANOVA. A preprint version of this manuscript is available on a preprint server (BioRxiv). The data that supports the findings of this study are available from the corresponding author upon reasonable request.

## Notes

### Competing Interest Statement

R. Polt, M. L. Heien and T. Falk have a financial interest in Teleport Pharmaceuticals, which expects to license the PACAP glycosides from Tech Launch Arizona, the patent holder. The other authors have no competing interests to declare.

### Summary of Updates

Key Revisions in this Resubmission: Expanded Microdialysis PK Dataset (n=4): Increased the sample size for our in vivo microdialysis pharmacokinetic studies to n=4 per group. Using a time-locked, ratio-paired t-test, we confirmed a statistically robust increase in central exposure for TES2320 compared to its unglycosylated parent compound. Reframed PAC1R Extracellular Domain (ECD) Mechanics: Clarified the binding rationale for TES2320 to emphasize how C-terminal glycosylation modulates membrane partitioning and extracellular orientation rather than implying total ECD independence, aligning with established Class B GPCR dynamics. Detailed Biophysical & Neurocircuitry Mechanisms: Expanded the Discussion to address the dissociation between I_h currents and rebound bursting (implicating downstream T-type 〖"Ca" 〗^(2+) channels and NMDARs), postsynaptic retrograde signaling (NO), and local circuit nuances of glutamate and PACAP co-release. Methodological & Visual Clarifications: Updated Figure 3 with representative RNAscope micrographs, added visual stimulus waveforms to electrophysiology figures, and fully detailed our automated QuPath transcript quantification workflows in the Methods section. Contextualized Study Scope & Limitations: Expanded our Discussion regarding behavioral battery scope, dose-selection rationale based on prior preclinical stroke/TBI models, and future directions for evaluating sex differences.

